# Synaptic Self-Organization of Spatio-Temporal Pattern Selectivity

**DOI:** 10.1101/2021.11.12.468380

**Authors:** Mohammad Dehghani-Habibabadi, Klaus Richard Pawelzik

## Abstract

Spiking model neurons can be set up to respond selectively to specific spatio-temporal spike patterns by optimization of their input weights. It is unknown, however, if existing synaptic plasticity mechanisms can achieve this temporal mode of neuronal coding and computation. Here it is shown that changes of synaptic efficacies which tend to balance excitatory and inhibitory synaptic inputs can make neurons sensitive to particular input spike patterns. Simulations demonstrate that a combination of Hebbian mechanisms, hetero-synaptic plasticity and synaptic scaling is sufficient for self-organizing sensitivity for spatio-temporal spike patterns that repeat in the input. In networks inclusion of hetero-synaptic plasticity leads to specialization and faithful representation of pattern sequences by a group of target neurons. Pattern detection is robust against a range of distortions and noise. The proposed combination of Hebbian mechanisms, hetero-synaptic plasticity and synaptic scaling is found to protect the memories for specific patterns from being overwritten by ongoing learning during extended periods when the patterns are not present. This suggests a novel explanation for the long term robustness of memory traces despite ongoing activity with substantial synaptic plasticity. Taken together, our result promote the plausibility of precise temporal coding in the brain.

**Author summary:** Neurons communicate using action potentials, that are pulses localized in time. There is evidence that the exact timing of these so called spikes carries information. The hypothesis, however, that computations in brains are indeed based on precise patterns of spikes is debated, particularly because this would require the existence of suitable detectors. While theoretically, individual neurons can perform spike pattern detection when their input synapses are carefully adjusted, it is not known if existing synaptic plasticity mechanisms indeed support this coding principle. Here, a combination of basic but realistic mechanisms is demonstrated to self-organize the synaptic input efficacies such that individual neurons become detectors of patterns repeating in the input. The proposed combination of learning mechanisms yields a balance of excitation and inhibition similar to observations in cortex, robustness of detection against perturbations and noise, and persistence of memory against plasticity during ongoing activity without the learned patterns. The proposed learning mechanism enables groups of neurons to incrementally acquire sets of patterns thereby faithfully representing their ’which’ and ’when’ in sequences. These results suggest that computations based on spatio-temporal spike patterns might emerge without any supervision from the synaptic plasticity mechanisms present in the brain.

## Introduction

Despite decades of research, it is still debated which coding schemes are used in central nervous systems. While in early sensory areas of cortex, stimuli appear to be represented mostly by spike rates, it cannot be disputed that temporal information is faithfully processed. While this can in principle be achieved by modulated spike rates in large populations of neurons it is tempting to hypothesize that at least in higher areas, as, i.e., frontal cortex, temporally precise responses of individual neurons play an important role. In fact, experimental studies on visual, auditory, olfactory, and somato-sensory cortex indicate that neurons can respond rather deterministically to inputs, underlining the possibility of precise spike codes.^1–8^

A range of theoretical studies attempted to elucidate mechanisms that could support precise coding of spatio-temporal patterns^9, 10^. It was found that with suitable synaptic weights, even simple integrate-and-fire neurons are sensitive to specific spatio-temporal input spike patterns. For instance, the Tempotron ^9^ was introduced as an extension of the Perceptron ^11^ to perform classification and detection of spatio-temporal patterns with a spike response to patterns only from a given set with a supervised algorithm for potentiating and depressing a neuron’s afferents. The number of patterns that a neuron can learn to classify depends on their length, the time constants of the neurons and the synaptic inputs ^12^. While in the Tempotron the action potential is allowed to occur anywhere during the time of the learned patterns, it was later shown that neurons can be forced to fire also at a specific time ^10^ during a specific pattern’s presence, which can be achieved by several more or less realistic synaptic mechanisms ^13, 14^. Both the Tempotron and the Chronotron employ supervised learning mechanisms based on label and time, respectively.

Supervised learning of spatio-temporal patterns seems at odds with reality, where the input is not labeled, impinges on the neuron continuously, and is subject to distortions and noise. In particular, it would need to explain how synaptic plasticity mechanisms become informed which aspect of the data should be taken into account when a label comes only long after the patterns. A recent study addressed this latter problem. It showed that neurons can recognize spatio-temporal patterns embedded in a background of noise using only weak supervision where the known number of repetitions of a pattern is used for optimizing synaptic efficacy ^15^. Based on the N-methyl-D-aspartate (NMDA) receptor^16–18^, a learning rule was proposed ^15^ that yields similar results as obtained by optimization. The biological plausibility of this correlation-based rule, however, is questionable. It does not respect Dale’s rule since synapses can change their sign, it is still supervised, and it requires a careful selection of potential weight changes such that only a given small percentage of potential changes become effective.

Therefore, it remains an open question if existing mechanisms of synaptic plasticity can be identified which enable neurons to specialize on statistically dominant patterns in the temporal stream of their inputs neurons in an entirely unsupervised manner.

The starting point of our considerations is synaptic scaling: It has been shown that neurons do not remain silent for long periods, but scale their weights to achieve a genetically intended number of spikes ^19^. Then, we use basic Hebbian mechanisms for the plasticity of both, excitatory and inhibitory neurons which for excitatory synapses resemble the NMDA-receptor. Dale’s law is enforced, i.e., inhibitory and excitatory neurons can not change into one another throughout the learning process. Furthermore, upper bounds on synaptic efficacies are imposed which prohibit the well-known runaway instability of Hebbian mechanisms. For excitatory synapses, we finally employ hetero-synaptic plasticity mechanisms that provide negative changes of efficacy, induce synaptic competition and thereby serve specialization^20^.

A combination of these realistic mechanisms turns out to be sufficient for the self-organization of pattern detection in single neurons. At times when no pattern is present excitatory and inhibitory inputs become globally balanced. During the time when a learned pattern is present we find detailed balance, where excitatory and inhibitory inputs cancel each other in temporal detail. These results parallel findings of the global and detailed balance of excitation and inhibition in cortex ^21–23^.

The resulting synaptic efficacies are then shown to ensure robust pattern recognition that is resistant to temporal jitter and noise. When basing learning on jittered patterns and also on Poisson rate modulations instead of precisely repeating patterns we obtain even more robust pattern detection. These results underline that the proposed mechanisms can be based on imprecise patterns and temporally modulated rate codes.

Next, we wondered if and how learned memory traces vanish during ongoing plasticity when only random patterns are presented which contain no statistically dominant structures. We find extremely long memory persistence already for a single output neuron. This leads to the question if the proposed mechanisms might contribute to solving the stability-plasticity problem^24^, such that they would support incremental learning where patterns occur rarely and are intermingled with random activity and/or different patterns. We investigated this for groups of output neurons where competition for patterns is induced by pre-synaptic hetero-synaptic plasticity ^25^. Thereby the output neurons specialize on different subsets of patterns such that the group as a whole self-organizes faithful representation of the ‘which’ and ‘when’ of patterns in the input. The memory persistence in this setting is finally shown to allow for incremental learning of sets of patterns in neuronal populations.

## Results

In all simulations, we consider simple leaky integrate and fire neurons ^26, 27^ with a fixed membrane time constant of 15 ms and pre-synaptic spikes originating from 500 input neurons (80% excitatory and 20% inhibitory). They provide input currents via kernels that have the shape of alpha-functions. For each synapse, the amplitudes of the input currents depend on a single parameter, the synaptic weight. The kernels have different time constants for excitatory and inhibitory synapses. The signs of the weights are not allowed to change during learning, i.e., Dale’s law is enforced.

### Single post-synaptic neuron

Synaptic plasticity is based on correlations between the input-kernels and deflections of the membrane potentials with respect to a threshold. For changes of the inhibitory synapses, positive deflections increase the weights, and negative deflections lead to their decay. For changes of the excitatory synapses, we let only positive deflections contribute, mimicking NMDA-dependent mechanisms. Without further constraints, this latter Hebbian mechanism is unstable.

Runaway instabilities are avoided by a combination of three simple but biologically highly plausible mechanisms: First, unbounded growth is made impossible by clipping the weights at upper limits for excitatory and inhibitory synapses. Second, positive weight changes for excitatory synapses are quenched when the long time activity exceeds a pre-determined rate, which mimics synaptic scaling ^19^. Then no positive weight changes can take place and only the signals for decreasing excitatory weights come into effect. Third, negative weight changes for excitatory synapses are induced by hetero-synaptic plasticity. Specifically, we include post-synaptic hetero-synaptic plasticity where the potential weight changes of different afferents are made dependent such that the resources needed for strong increases of the post-synaptic contributions to synaptic efficacies are taken from synapses which would otherwise increase only weakly. Thereby the latter synapses’ efficacies become reduced. Note, however, that this does not imply strict normalization of excitatory weights (see Materials and Methods).

It turns out that these ingredients are sufficient for robust self-organization of spatio-temporal spike pattern detection in single neurons. As an example, Fig 1 shows the membrane potential (MP) of a single neuron before and after learning a random pattern of length 50 ms that has the same statistics as the random background but repeats in every training epoch of length 2000 ms.

**Fig 1.**
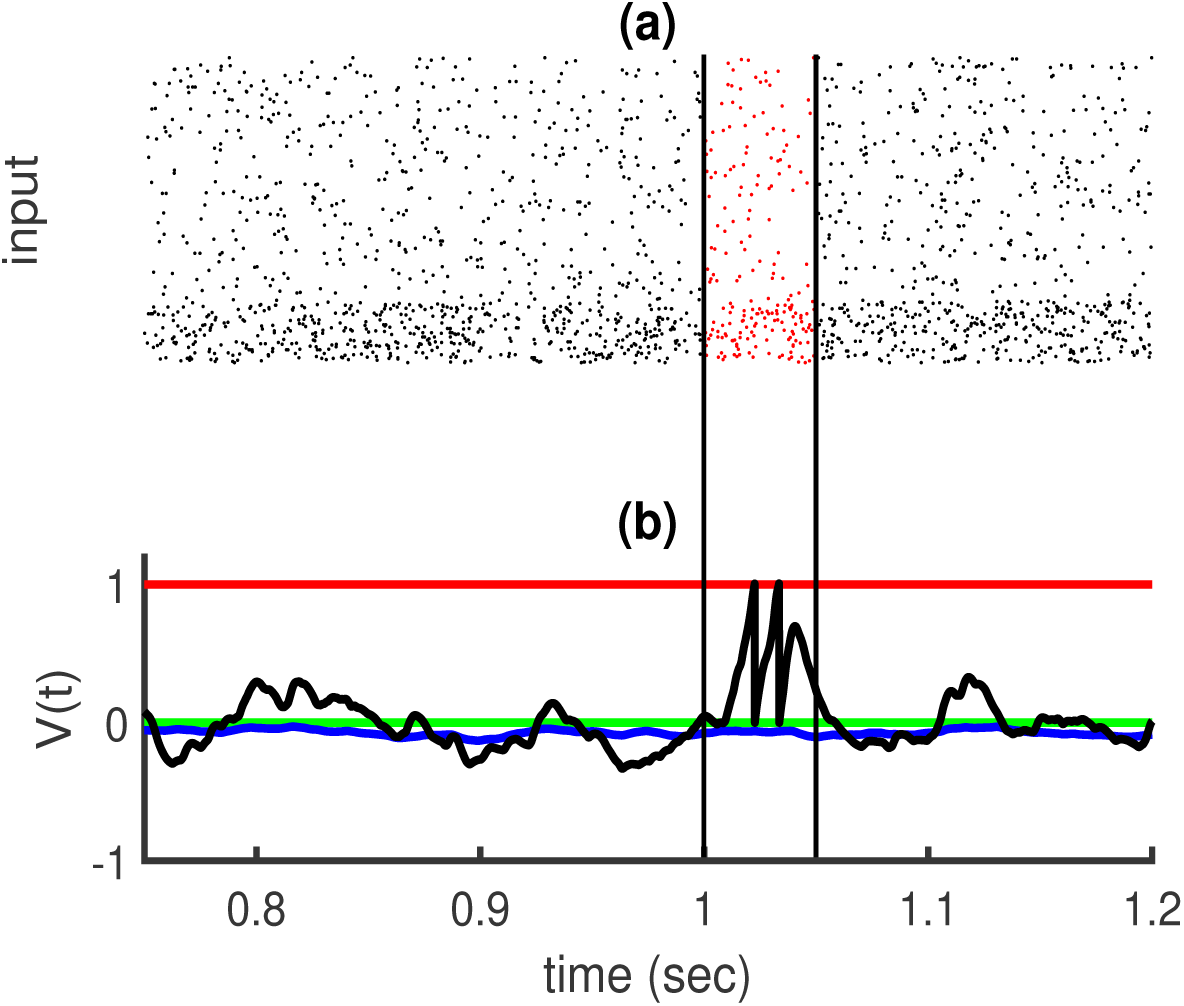
a: Input activity in the raster plot, five hundred afferents (80% excitatory and 20% inhibitory) send inputs to one post-synaptic neuron. The excitatory neurons’ rate is 5 Hz, and the inhibitory neurons’ rates are 20 Hz. There is a random embedded pattern of length 50 ms in the red area between black vertical lines. b: Membrane potential versus time. The black trace shows that after learning fluctuations outside of the embedded pattern are attenuated, and two spikes occur at the embedded pattern time (vertical black lines). The green line is for resting potential, red is for threshold, blue is for before learning, and black is for after learning.

After convergence of the weights, we separated the excitatory and inhibitory inputs to see how their respective contributions lead to firing only during the pattern (Fig 2 a).

**Fig 2.**
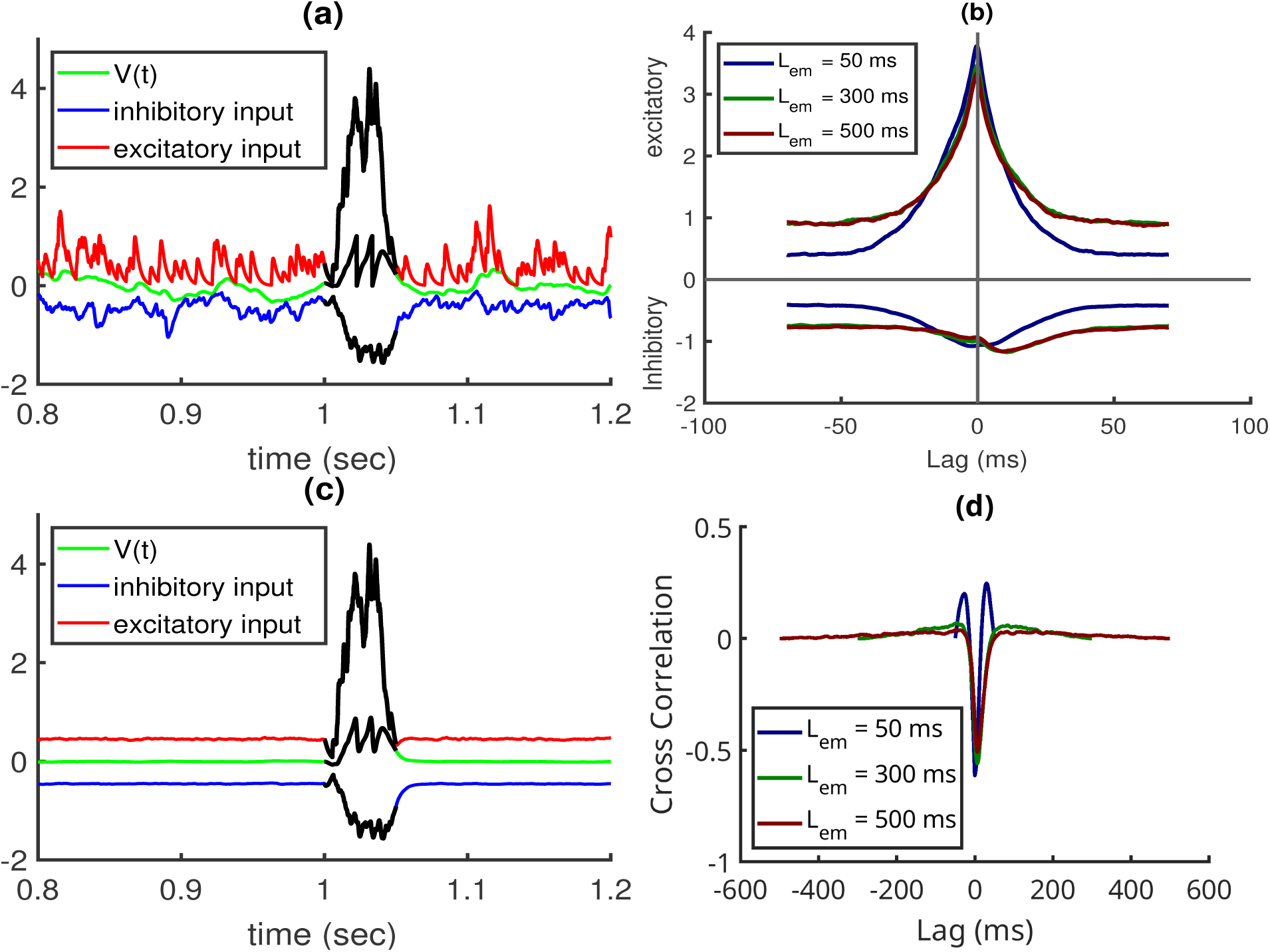
a,b,c: A fixed pattern is embedded between 1000 ms and 1050 ms, i.e. duration *L*_*em*_ = 50ms. a: excitatory and inhibitory inputs and the membrane potential after convergence of self-organization with this pattern. b: Spike triggered average for different pattern lengths. The inhibition minimum follows the maximum of excitation in the more extended pattern. c: Average contribution of excitatory and inhibitory inputs from 1000 epochs with different epochs. d: cross-correlation between inhibitory and excitatory inputs for neurons that where exposed to patterns of *L*_*em*_ = 50, 300, and 500 ms (shown in their afferents starting from 1000 ms.). The shown correlations are averages from 1000 simulations with random patterns. In all figures, the length of the training epoch is 2000 ms, and the desired number of spikes is 2, i.e., the desired firing rate is *r*_0_ = 1 Hz.

Fig 2 a indicates that inhibitory and excitatory inputs cancel each other outside of the embedded pattern in the mean (global balance). Fig 2 b depicts the spike-triggered average of the inhibitory and excitatory input, respectively. Averaging these inputs on epochs (Fig 2 c) explicitly shows that global balance removes fluctuations in the mean. In contrast, during the time of the embedded pattern, the respective contributions to the membrane potential both increase but remain mostly balanced also across time (detailed balance). In fact, only some residual imbalance between excitatory and inhibitory afferents allow the post-synaptic neuron to fire during this period of time. The detailed balance during the embedded pattern is reflected by a substantial anti-correlation between excitatory and inhibitory afferents at zero time shift (Fig 2 d).

To quantify the performance of the learning mechanism for ensembles of random patterns, we consider the average percentage *R* of spikes that correctly detect the pattern:

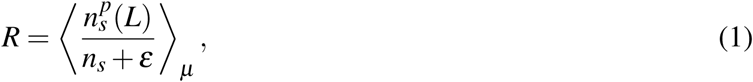

where *n*_*s*_ is the total number of spikes during a testing period and 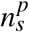 the spikes’ number related to the presence of the pattern to be detected. Since patterns can induce spikes also shortly after the pattern due to the finite decay time of the excitatory synaptic kernel, the time window for testing if spikes occur inside the pattern is extended by *L* ms after the pattern has ended. Adding an arbitrary small number *ε* to the denominator ensures a definite result (*R* = 0) when no spikes occur at all. The ratio is averaged for an ensemble of independent embedded patterns *µ*.

Occasionally, we also consider a variant of this criterion *R*\* where the average is taken only on the epochs in which there is at least one spike occurring in post-synaptic neuron:

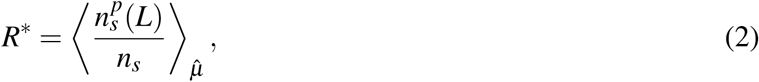

where 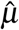 refers to the ensemble of independent embedded patterns for which the post-synaptic neuron elicited at least one spike during the pattern presentation.

Fig 3a shows that neurons learn to fire only in response to the embedded pattern and remain silent otherwise. Extending the testing window by *L* = 15 ms reveals that the performance becomes perfect. Fig 3b shows that using longer embedded patterns allows neurons to learn them faster: a more extended embedded pattern has more space for residual excitatory-inhibitory imbalances and more contribution to the weight changes; therefore, it can find the embedded pattern more rapidly.

**Fig 3.**
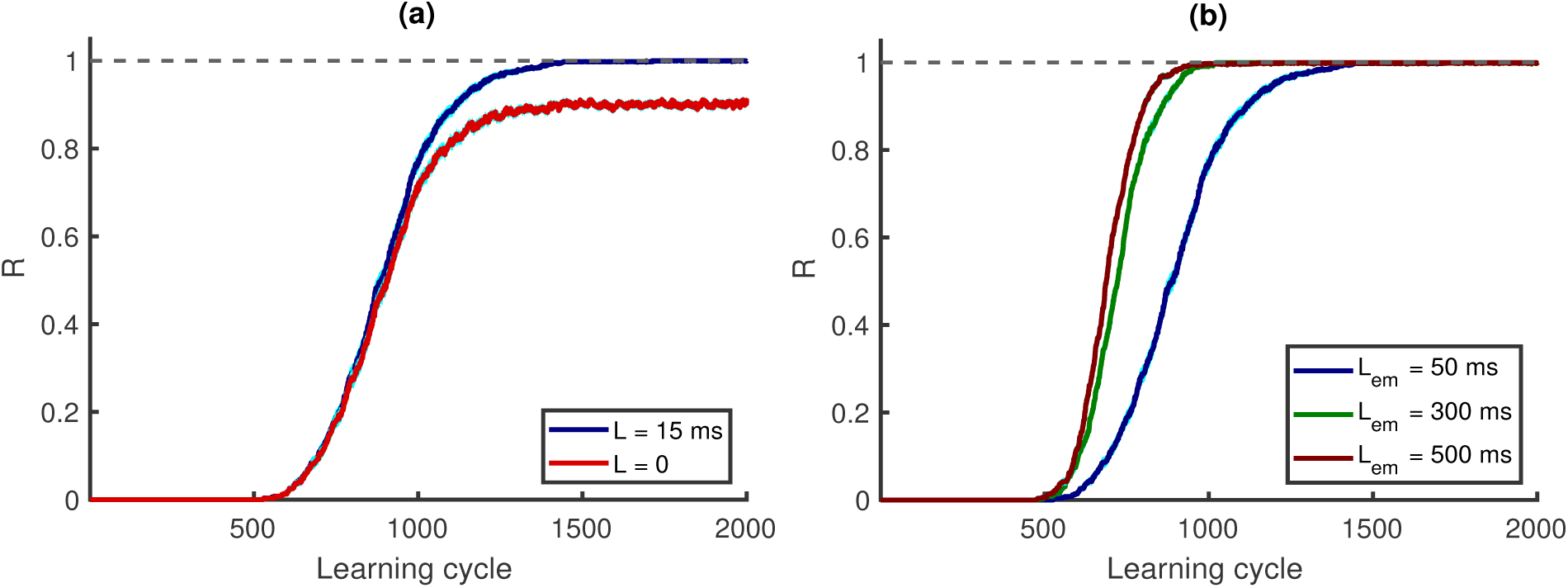
(Color online) a: Learning performance R versus learning cycle, for *L* = 0 and *L* = 15 ms. A 50 ms pattern is embedded between 1000 and 1050 ms. b: R versus learning cycle, *L* = 15 ms. Patterns duration,*L*_*em*_, are 50, 300, and 500 ms, shown in afferents from 1000 ms. R is an average of 500 simulations, in which there are 500 afferents, and the length of the training epoch is 2000 ms. The desired firing rate is *r*_0_ = 1 Hz.

### Noise and memory robustness

We wondered if and how the memory for the originally learned pattern decays when synaptic plasticity is present during long periods of random inputs where an already learned pattern does not re-appear. Note that plasticity was switched off when we tested whether it remembers the originally learned pattern. Fig 4 a shows that even after 20000 learning cycles, 80 percent of spikes would still occur during the embedded pattern time, (chance level is 0.025). In particular, after a dropping to this value *R* remains constant for a period that would correspond to more than 10 hours, with practically no further decay. This striking memory persistence can be understood by considering that due to the synaptic scaling inherent in the learning mechanism, the neurons will change synapses until the pre-determined long-term firing rate is achieved also for random patterns (red in Fig 4 b). Since the inputs have no structure, the weights mostly become only scaled, which per se cannot erase the selectivity for the learned pattern. When the learned pattern is then again presented to the afferents, the neuron fires additional spikes during the pattern, which leads to a much higher firing rate for the pattern (green in Fig 4 b).

**Fig 4.**
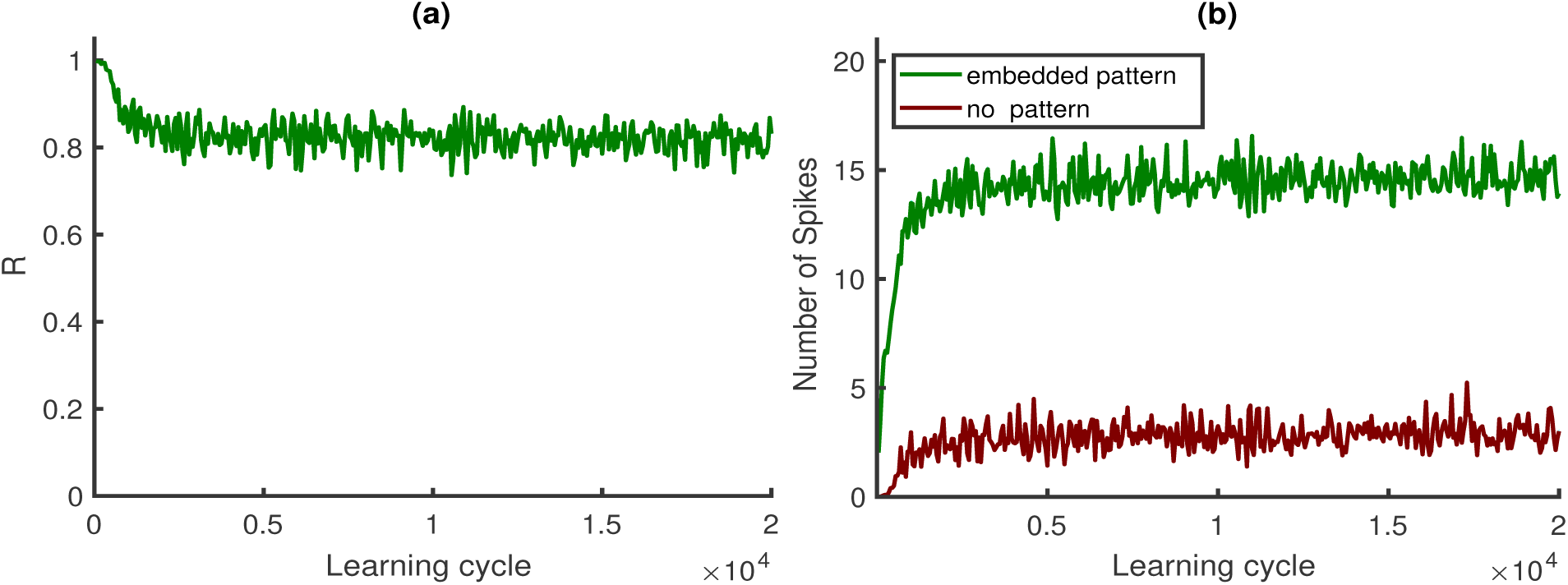
(Color online) a: *R* as memory criterion versus learning cycle: Without learned patterns in afferents, learning continues, and every 50 cycles, learning is paused, and the learned pattern is placed in the background, then R is computed as a memory criterion. b: number of spikes versus learning cycle. Neuron elicits more spikes when the embedded pattern is in the afferents (green) than the situation without a pattern(red). Patterns duration is 50 ms, shown in afferents starting from 1000 ms, *L* = 15 ms, and this figure is based on an average of 500 simulations.

The changes between a new weight vector and the initial weight vector can be in amplitude and angle. In order to keep the memory, the learned weight vector should not change its direction in weight space. If the cosine of the angle remains close to one, the neuron would still remember the pattern, while the change in the number of spikes is caused by increasing the norm of a weight vector. As Fig 5 a shows that the norm of the weight vector indeed changes dramatically while the angle changes are rather insignificant (Fig 5b).

**Fig 5.**
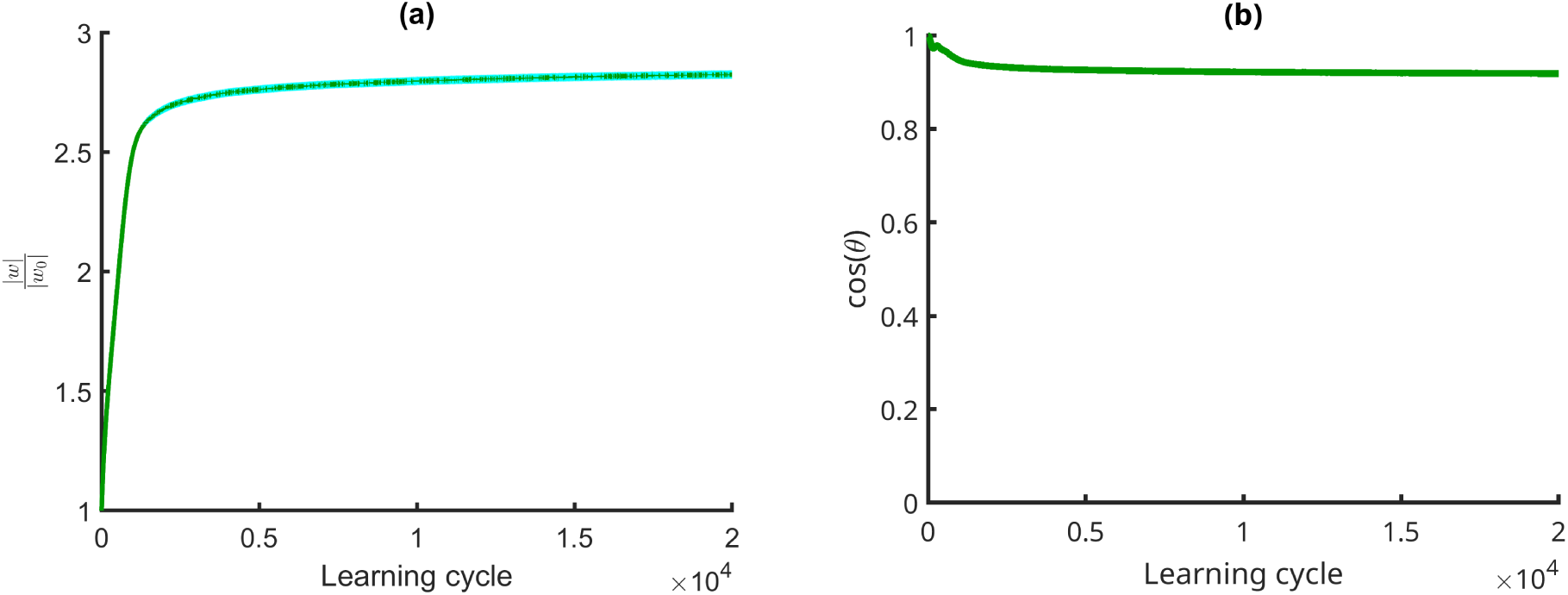
(Color online) Without learned patterns in afferents, learning continues. a: norm of the current weight vector divided by the learned pattern weight vector (*w*_0_). b: the cosine between the current weight vector and *w*_0_. This figure is based on an average of 500 simulations.

Taken together, we find that when learning is continued with random input the neuron achieves the genetic firing rate mostly by scaling its synapses up and then randomly fires in response to the noisy input.

Taking these findings into account both learning and persistence of pattern selectivity can heuristically be understood in combination. Let’s first consider the weight changes caused by stochastic background only. Here, the instability of Hebbian excitatory plasticity drives a subset of weights to large values until the desired number of spikes occurs in the mean. Simultaneous Hebbian plasticity of inhibition ensures that finally global balance is achieved. Then, the neuron is in the fluctuation driven regime, with rather strong excitatory and inhibitory weights. After achieving this balance further weight changes induced by the stochastic background induce mainly some random walk confined around this fixed point.

This raises the question if the fixed point of the weights learned from random input alone is too stable to allow for subsequently learning a particular pattern. Fig 6 demonstrates that learning is indeed possible also with this initialization. Intuitively, the weights perform a random walk around the fixed point when only random input is presented, but when a repeating pattern is introduced an additional drift systematically shifts the weights away from this fixed point until the desired number of spikes occur only for the pattern and none in response to the background ^28^. Note, that this entails that some of the weights originally acquired from the background decay. This weight dynamics continues until the spikes occur only in the period of the repeating pattern. Then the remaining weight changes cause diffusion around this new fixed point. Hence, also with such an initialization the neuron can learn to fire in response to the embedded pattern. The learning speed, however, is reduced. Fig 6c shows that the R-value approaches one, and Fig 6b that the cosine between the new weight vector and the initial weigh vector changes.

**Fig 6.**
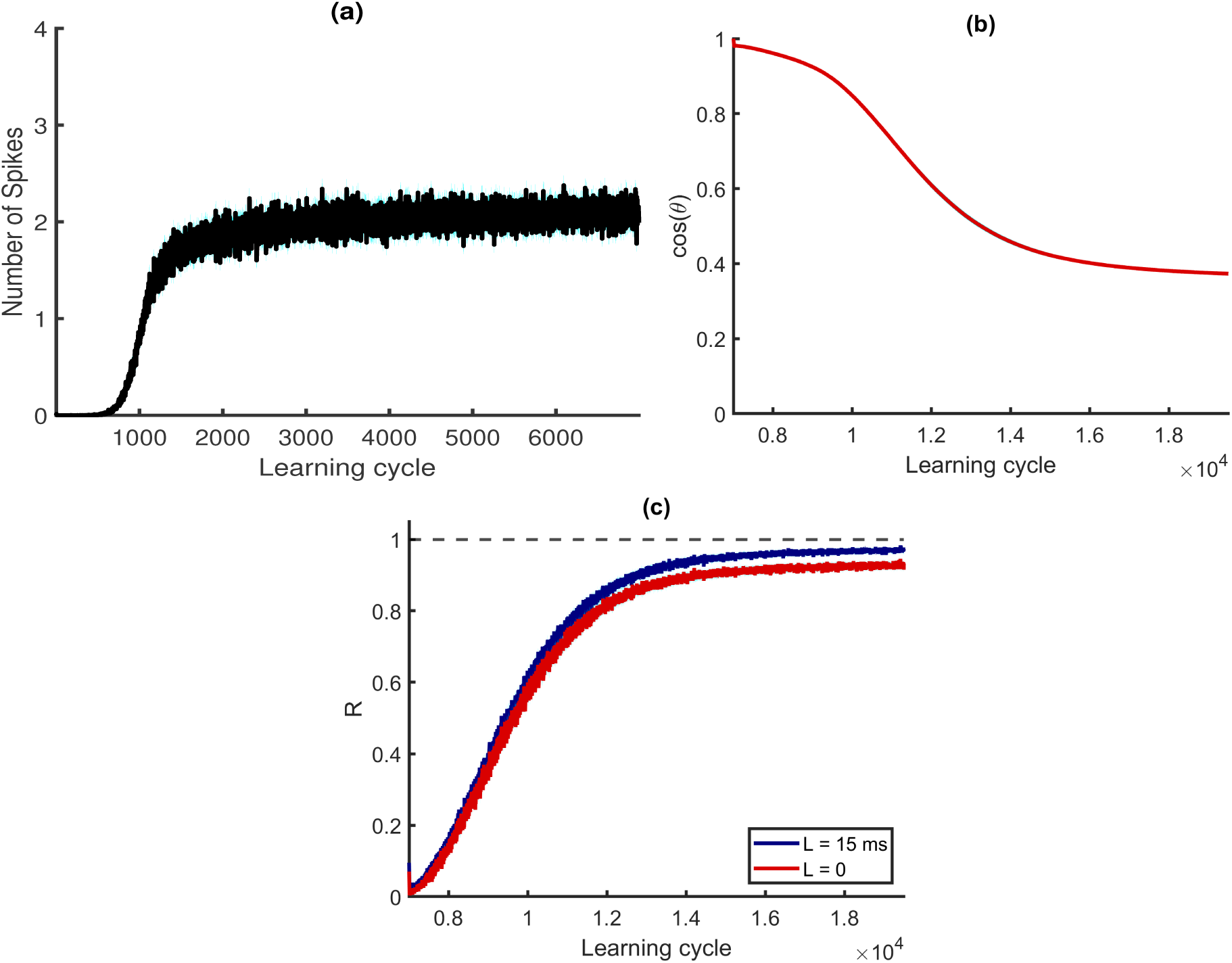
(Color online) There is no embedded pattern till learning cycle 7000 in afferents, and there is a 50 ms embedded pattern in afferent from learning cycle 7001 shown in afferents from 500 ms. a: number of elicited spikes versus learning cycle, the post-synaptic neuron fires two times in response to the noisy background. b: the cosine between the weight vector and the initial weight vector at learning cycle 7000. c: Learning performance R versus learning cycle, for *L* = 0 and *L* = 15 ms. This figure is based on an average of 500 simulations, in which there are 500 afferents. The length of the training epoch is 800 ms, and the desired firing rate is *r*_0_ = 2.5 Hz. For more details see Materials and Methods.

In order to understand why the memory for the learned pattern persists when learning continues without the embedded pattern being present note that the weights for pattern detection are specific for the repeating pattern and at the same time guarantee that the background alone will not elicit spikes. When then learning continues without the repeating pattern all weights are mainly scaled up due to the instability of the Hebbian mechanism for excitatory synapses. In consequence the norm of the weight vector grows (Fig 5 a) while its direction is preserved (Fig. 5 b). Thereby the structure of the learned weights persists. A more detailed mathematical analysis of this striking persistence of the memory for spatio-temporal patterns is in preparation, however, goes beyond the scope of this paper.

The plausibility of spike pattern coding depends on its robustness against noise and pattern distortions. For neurons that have learned a particular pattern without noise we examined the dependency of detection performance on three types of noises:

**First**: we removed spikes inside the embedded patterns and examined if neurons can still recognize the embedded pattern. Fig 7a shows the robustness against removing spikes inside the embedded pattern.

**Fig 7.**
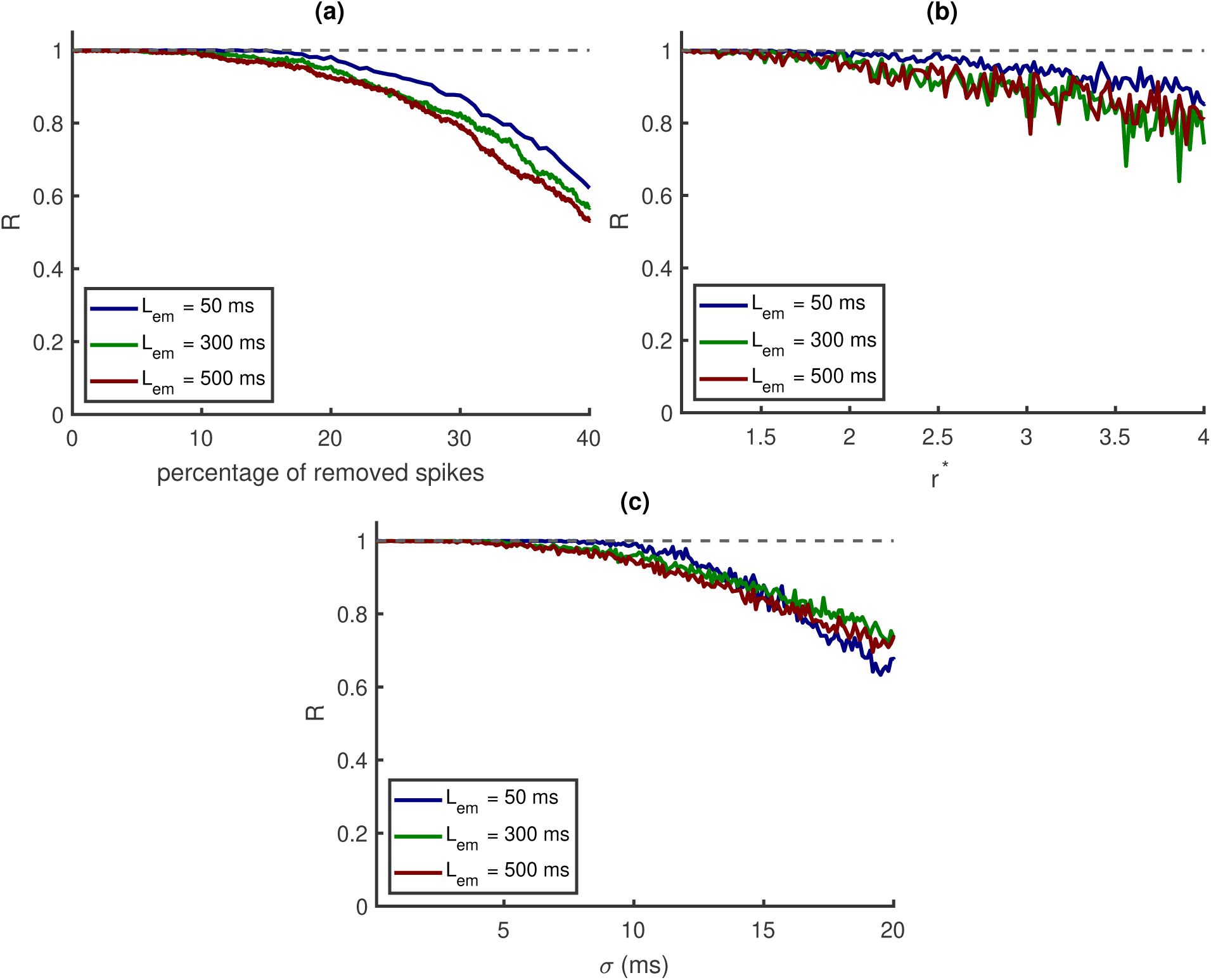
Effect of noise on the learning algorithm. a: removing spikes. b: increasing firing rates. c: Jitter noise. Patterns durations are 50, 300, and 500 ms, shown in afferents starting from 1000 milliseconds. This figure is based on an average of 500 simulations, in which there are 500 afferents, the length of the training epoch is 2000 ms, and *r*_0_ = 1 Hz.

**Second**: during learning, the neuron receives inhibitory input at a frequency of 20 Hz and excitatory input at a frequency of 5 Hz (ratio 4 to 1). For testing we added *S* random spikes per second to excitatory input neurons and *4* × *S* spikes per second into inhibitory neurons. The new rates becomes 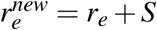 and 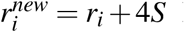. Fig 7b shows that the performance R decays when 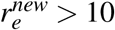. Here we define 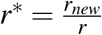.

**Third**: in this part, we examine the robustness of detection against jitter noise. For this purpose, we shuffle the times of the spikes in the afferents using a Gaussian distribution with zero mean and *σ* standard deviation. Fig 7c shows that the algorithm is robust until *σ* = 5 ms, and after that, performance starts to decay.

Next, we wondered if precise patterns are required for self-organizing pattern selectivity. First, we perturbed the training patterns by jittering the spikes according to a Gaussian distribution with mean zero and a standard deviation of 20 ms. We found that this does not hamper learning. Fig 8a shows the R-value in each learning cycle; the blue line is for testing with the original pattern, the green line with the jittered patterns. The red line represents *R*\*, i.e. we dropped contributions to R from the epochs where no spikes at all occur. Then, we converted each afferent’s time code input to rate code input (i.e. Poisson spike rates). For this purpose we first convolve each spike with a Gaussian distribution with a zero mean and a standard deviation of 20 ms. The resulting function is then used as modulated firing rate of a Poisson point process. This results in an ensemble of spatio-temporal patterns based on the original pattern that has the statistics of Poisson processes, including failures and a Fano factor of 1. These distorted patterns are then used for learning and testing. This transformation is leading to a similar result (Fig 8b, the blue line is for testing with the original pattern, the green line for testing with Poisson spike rates including no spikes, the red when epochs with no spikes are dropped).

**Fig 8.**
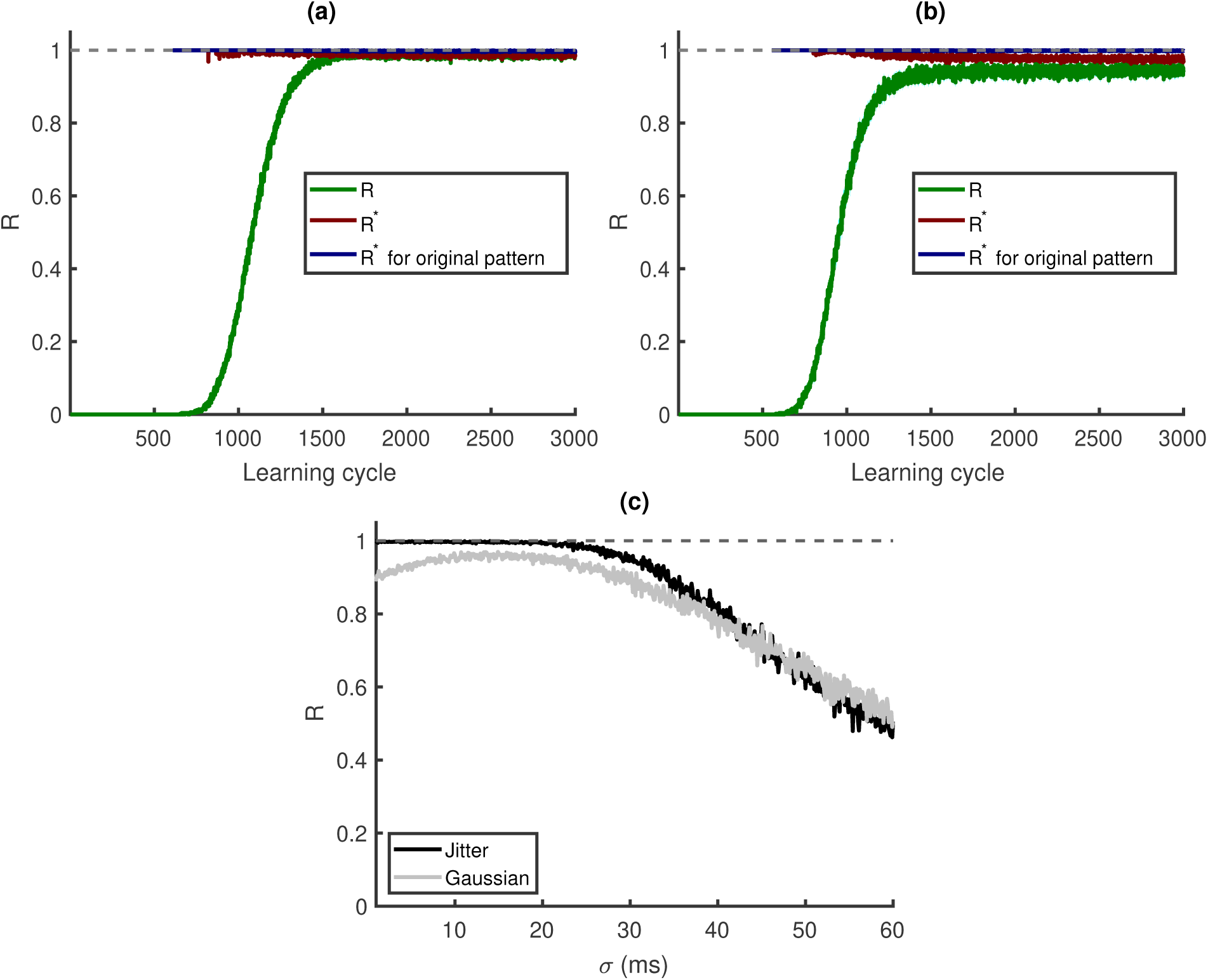
Learning with jittered patterns and Poisson spike rates: a: the training patterns are perturbed by jittering the spikes with a Gaussian distribution with a standard deviation of *σ* = 20 ms and a zero mean. Blue represents testing with the original pattern (*R*), the green line represents jittered data with no spikes in MP (*R*), and the red line represents dropping episodes with no spikes in MP (*R*\*). b: Poisson spike rate learning: the blue line represents testing with the original pattern (*R*\*), the green line represents testing with Poisson spike rates, including those with no MP spikes (*R*), and the red line shows *R*\* when dropping episodes with no spikes. c: the black and gray lines represent tests of jittered and Poisson spike rates, respectively, with varying *σ*. This figure is based on an average of 500 simulations with 500 afferents. The training epoch is 2000 ms, *L*_*em*_ = 50 ms, *L* = 15 ms, and *r*_0_ = 1 Hz.

It turns out that with the corresponding weights pattern detection becomes more robust with respect to jitter noise (Fig 8c, the blue and red lines are for testing with different *σ* for jittered and Poisson spike rates, respectively). These results demonstrate that codes based on temporal rate modulations and spatio-temporal pattern detection by individual neurons are compatible.

### Diversification by pre-synaptic hetero-synaptic plasticity

In the approach presented so far a neuron can become a detector for more than only one pattern, along the lines of the Tempotron ^9^ (Fig 9). Its activity would then, however, obscure which individual pattern was present at which time. For a faithful representation of a sequence of patterns, as, e.g., the phonemes in spoken human language, it is therefore desirable that different neurons in a network become specialized for different subsets of patterns such that as a whole a network represents precisely the which and when of patterns in a sequence. If successful, such a system could, e.g., be used for unsupervised speech recognition ^15^.

**Fig 9.**
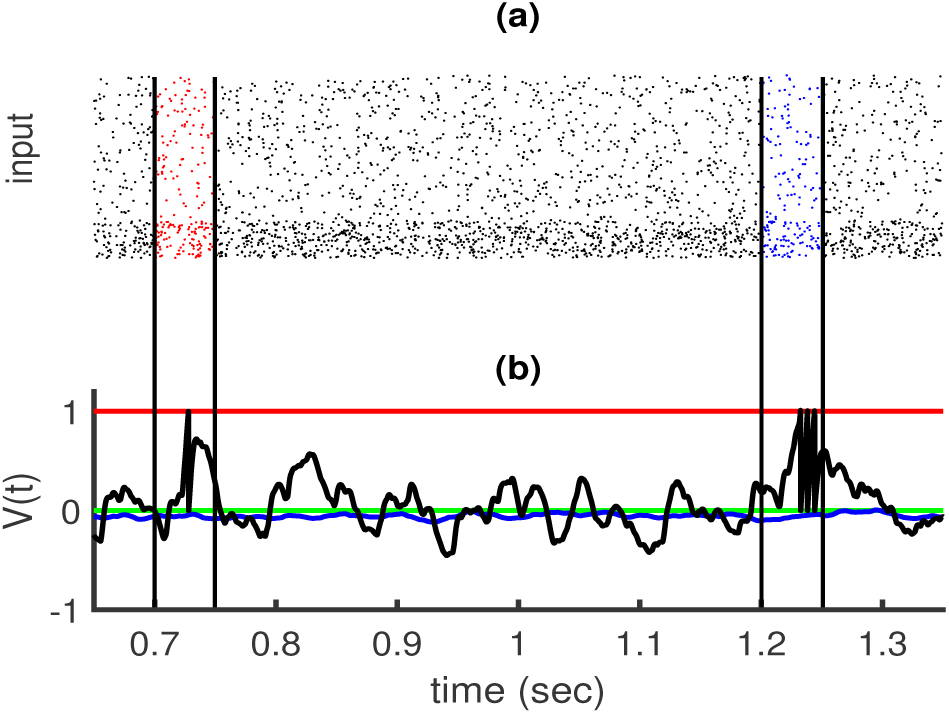
Example of membrane potential versus time after learning with two patterns: a: Input activity in the raster plot, five hundred afferents (80% excitatory and 20% inhibitory) send inputs to one post-synaptic neuron. The excitatory neurons’ rate is 5 Hz, and the inhibitory neurons’ rates are 20 Hz. There are two random embedded patterns of length 50 ms in the red and blue areas (between black vertical lines), *r*_0_ = 2 Hz, and the length of the training epoch is 2000 ms. b: Membrane potential versus time. The black trace shows that after learning neuron responds to both embedded patterns. The green line is for resting potential, red is for threshold, blue is for before learning, and black is for after learning.

As a first step into this direction we wondered which realistic synaptic mechanism might enforce the specialization of different neurons in a network for different pattern sets. We found that already synaptic competition induced by pre-synaptic hetero-synaptic plasticity^25^ yields sufficient selectivity for different patterns in an ensemble of neurons such that identity and order of patterns are represented faithfully (for details of the computational implementation see Materials and Methods).

To quantify the performance, we use the rank of a matrix where each row represents one of the neurons, and each column one of the patterns. The matrix element in each row is set to 1 if the corresponding neuron is active for that pattern, and to 0 otherwise. The rank of this matrix provides the number of linearly independent row vectors. When it meets the number of the different patterns in a given stimulus the population of neurons faithfully represents the presence and the order of the patterns. Therefore, we introduce the ratio of the rank of this matrix to the number of patterns as performance measure Ω.

As an example, we trained networks of different sizes with four patterns where always all patterns were present in each training epoch. We found that the pre-synaptic competition leads to selectivity for all four patterns as soon as 14 post-synaptic neurons were present. In contrast, without this pre-synaptic competition, separation does not become complete.(Fig 10 a).

**Fig 10.**
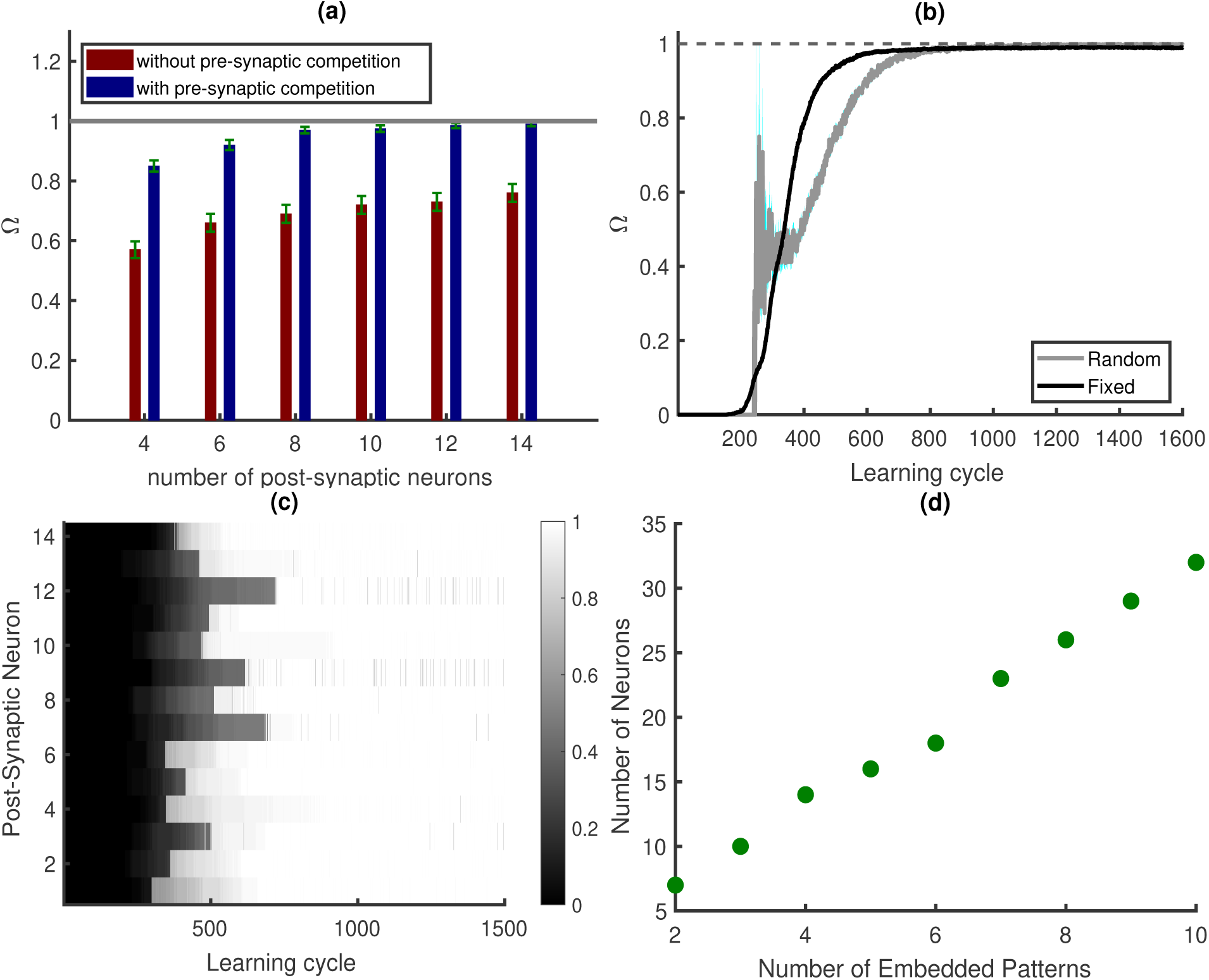
a: The system’s Ω (rank/4) versus the number of post-synaptic neurons after learning. b: Ω versus learning cycle (There are 14 post-synaptic neurons). Black line: Each pattern is shown in each learning cycle in a fixed position. Gray line: Each pattern is shown in the epoch with the probability of 0.2 at a random place. c: Each row shows the mean R values for the case of 14 post-synaptic neurons. d: number of postsynaptic neurons to separate different numbers of embedded patterns. a, c, and d are based on an average of 50 simulations, and b is based on an average of 500 simulations. In all simulations, the epoch length is 2000 ms, and there are 4 independent embedded patterns, with a duration of 50 ms,*L* = 15 ms, and *r*_0_ = 1 Hz.

We then tested the persistence of learned selectivities (with 14 neurons) when learning is continued with random input. We tested performance (with plasticity switched off) and found practically no decay of Ω even when learning was continued 10X longer than it takes for learning all patterns.

This result motivated us to test if learning a set of patterns is possible from learning epochs that contain only subsets of all patterns. Our preliminary investigations indicate that such incremental learning is indeed possible. As an example, we considered 4 embedded patterns and 14 post-synaptic neurons. Each embedded pattern has a 0.2 probability of being present in each learning cycle in the epoch. Note that, thereby some learning cycles have no embedded pattern in the epoch. Also, we randomly chose locations from ([300 300 + *L*_*em*_), [400 400 + *L*_*em*_), …[1700 1700 + *L*_*em*_] ms) to put the embedded patterns in them.

As Fig 10b shows the relative rank Ω goes to one, indicating that patterns can be learned also incrementally and R value goes to one for all postsynaptic neurons Fig 10c.

We then test how many post-synaptic neurons are required to separate different numbers of embedded patterns. Fig 10d shows a linear increase in the number of post-synaptic neurons necessary for selecting different numbers of embedded patterns (where Ω⩾ 0.97).

## Discussion

Neurons respond faithfully to input sequences ^29^. That is, they are rather deterministic devices and with suitable synaptic efficacies, individual neurons can, in principle, serve as detectors for specific spatio-temporal input spike patterns. This opens the possibility that coding and computation in brains are at least in part based on temporally precise action potentials.

In the past, this hypothesis has been investigated in simple integrate and fire models. Supervised learning rules were proposed that enable neurons to signal the presence of a pattern ^9^ and to fire at predefined time points during a specific pattern ^10^. It was further demonstrated that relatively weak supervision can be sufficient for learning the synaptic weights for pattern detection ^15^. Here, only knowledge about the number of pattern occurrences is needed for the specialization of a neuron ^15^. While a learning rule for this ‘aggregate label learning’ was rigorously derived, the proposed biological realization suffers from several rather unrealistic assumptions. In particular, excitatory and inhibitory plasticity are not treated separately, and Dale’s law was not observed. When the signs of synapses may change, this has also a negative impact on the excitatory and inhibitory balance. For some patterns, there may be only few inhibitory neurons remaining in the system after learning. When there is a lack of inhibition; however, the potential can take a value close to the threshold, increasing the chance of getting random spikes outside the embedded patterns. Also, a selection criterion was used, by which independently of their sign, only the largest 10% of changes were taken into account, for which no realistic interpretation was provided.

Therefore, it remains an open question if the synaptic plasticity mechanisms present in real neuronal networks support a coding scheme that is based on spatio-temporal patterns of spikes.

We found that a combination of Hebbian mechanisms, hetero-synaptic competition, and synaptic scaling indeed makes individual neurons sensitive for statistically dominant spatio-temporal patterns in their afferents without any supervision.

Performance is found to be robust to temporal jitter, missing spikes, and additional noise. In particular, also patterns consisting of Poisson spike rate modulations are captured surprisingly well by the proposed plasticity mechanisms and lead to robust detection (Fig 7 b, c).

The proposed combination of learning mechanisms yields a detailed balance of excitation and inhibition where this is possible: during the learned pattern. This fits nicely to experimental observations ^21^ revealing a negative correlation between excitatory and inhibitory inputs. Outside the pattern, global balance is achieved ^23^.

Balance is a natural consequence of Hebbian mechanisms when they are simultaneously present in both excitatory and inhibitory synapses, and the otherwise unstable growth of excitatory efficacies is constrained. While this has been noted before ^30^, we here show that Hebbian plasticity can select synaptic efficacies that make neurons detectors for spatio-temporal patterns when realistic constraints are taken into account. In particular, we found that a subtle interplay of the instability of Hebbian mechanisms for excitatory synapses and synaptic scaling enforces a local imbalance during the learned patterns which leads to specific and temporally precise spike responses.

The excitatory-inhibitory balance protects memory for rate patterns in neural networks ^30, 31^. Our simulations now demonstrate that also the memory for spike patterns is protected from being overwritten by noise or other patterns.

After successful learning of patterns, the random background input does not lead to spikes in individual neurons. When then plasticity is continuously present for a long time during which only random inputs are present, the input weights mainly become scaled up until the desired number of spikes is reached, i.e., the plasticity mechanisms considered here are consistent with synaptic scaling ^19^. Obviously, scaling alone preserves the memory for the learned patterns, which will then induce far more spikes than the background. In other words, the plasticity mechanisms discussed here lead to a very long memory persistence already in a single neuron such that selectivity is preserved and sensitivity becomes even enhanced. We tested the memory persistence also in groups of neurons that specialized for subsets of the input patterns via pre-synaptic hetero-synaptic plasticity. Here, we observed practically no decay.

This finding suggests that in groups of neurons incremental learning of sets of patterns should be possible, wherein each successive training epoch only a subset or even none of all patterns is present. We could confirm this hypothesis for simple cases; however, the question if the combination of plasticity mechanisms discussed here indeed provides a solution for the notorious stability-plasticity dilemma ^24^ in spatio-temporal pattern learning will require more systematic investigations which go beyond the scope of this paper.

The fact that biologically realistic plasticity mechanisms can support the self-organization of spatio-temporal pattern detection by individual neurons underlines the possibility that such temporal codes are indeed present in nervous systems. Particularly the ability to learn patterns underlying Poisson spike rates demonstrates that such a temporal coding scheme can be consistent with rate codes. We speculate that this transformation of temporally modulated rates to spike pattern codes could explain the increase in sparsity observed in the early visual cortex when natural contexts are included ^32^ as well as the extreme sparsity of activations in higher cortical areas.

## Materials and Methods

### Neuron model

In this work, we employ the leaky integrate and fire (LIF) model. The dynamic of the membrane potential of a single neuron *V*(*t*) which is receiving current from *N* afferents is:

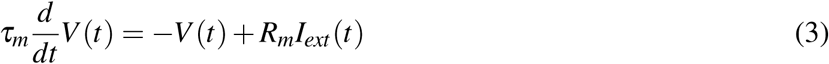

where *τ*_*m*_ and *R*_*m*_ are time constant and resistance of the membrane, respectively. We consider *R*_*m*_ = 1. Whenever the neuron arrives at or passes the threshold it evokes spike output and resets to resting potential. Resting potential is zero in this study, and *I*_*ext*_ is the external current from excitatory (E) and inhibitory (I) afferents:

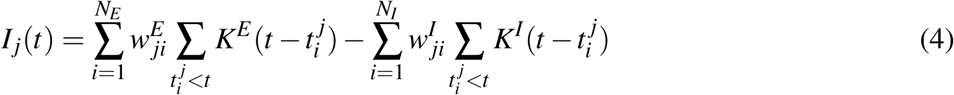

where 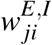 represents the respective *i*′*th* afferent’s excitatory and inhibitory synaptic strength to output neuron *j. N*_*E*_ and *N*_*I*_ are the numbers of excitatory and inhibitory afferents, respectively (*N* = *N*_*E*_ + *N*_*I*_). Note that at time 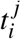, there is a spike in afferent *i*, and at this time, afferent *i* start to send input to the post-synaptic neuron *j* as much as *w*_*ji*_*× K*. The shape of the kernel is an alpha-function with the following equation:

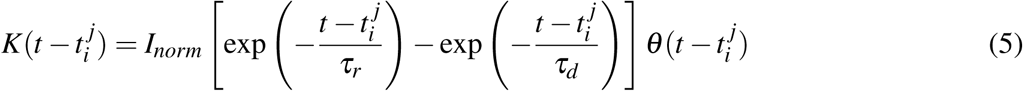

where *θ* is the Heaviside step function. *τ*_*r*_ and *τ*_*d*_ are time constants of synaptic current, which are different for excitatory and inhibitory synapses. In this study *τ*_*m*_ = 15 ms. 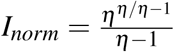 normalises *K* to unit amplitute, where 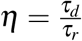. To numerically integrate equation 3, we use Euler method with Δ*t* = 0.1 ms. The parameters are found in Table 1.

**Tab 1.**
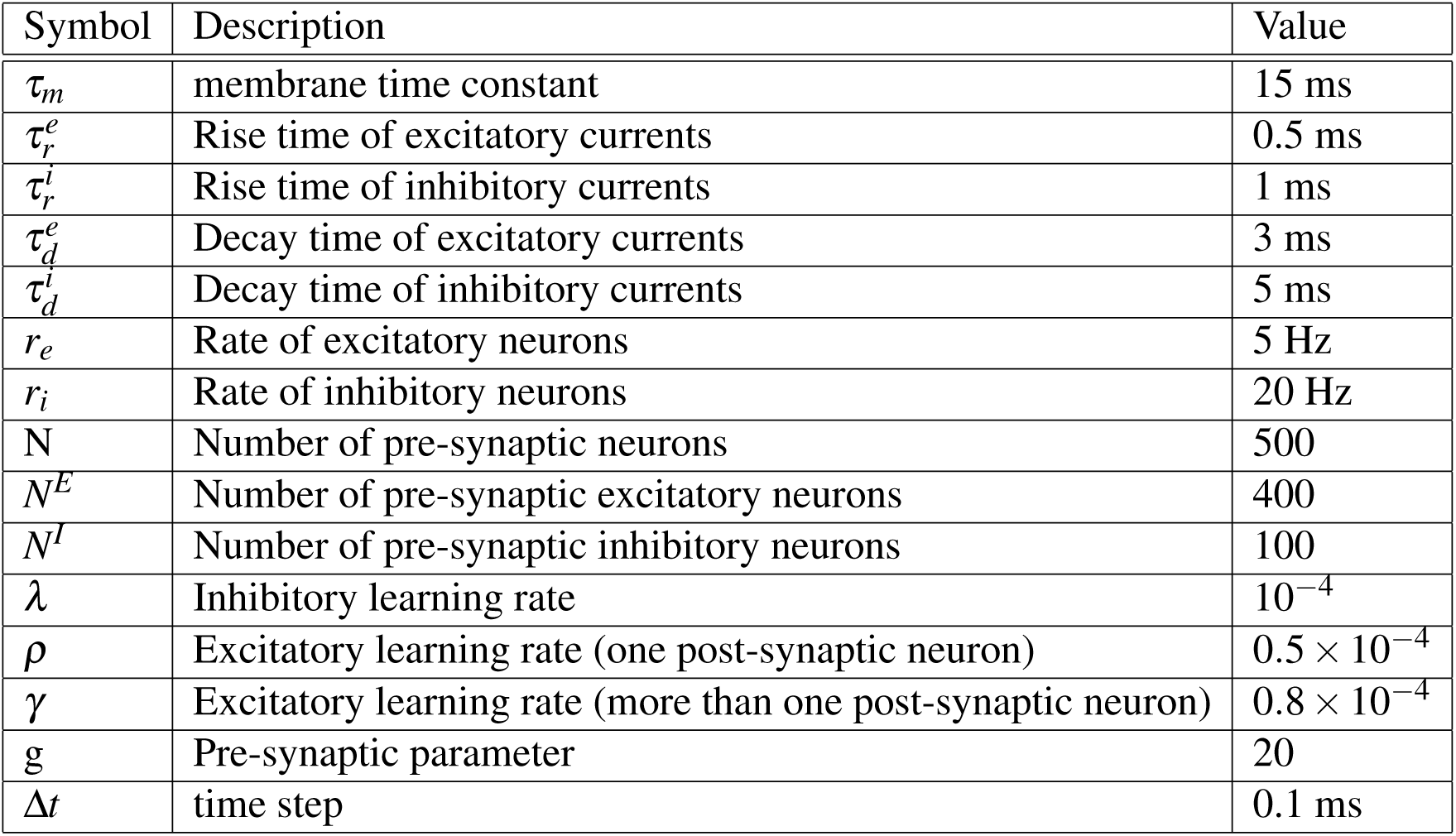
List of parameters.

### Learning algorithm

We assume that a synapse’s efficacy is the product of pre- and post-synaptic components.

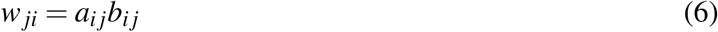

where *a*_*i j*_ is provided from the pre-synaptic neuron *i*, and *b*_*i j*_ is provided by the post-synaptic neuron *j* (Fig 11). If we have *N* pre-synaptic neurons (*i* = 1, …, *N*) and *M* post-synaptic neurons (*j* = 1, …, *M*) based on Taylor expansion small changes *δa*_*i j*_ and *δb*_*i j*_ lead to

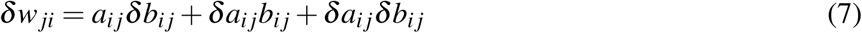

**Fig 11.**
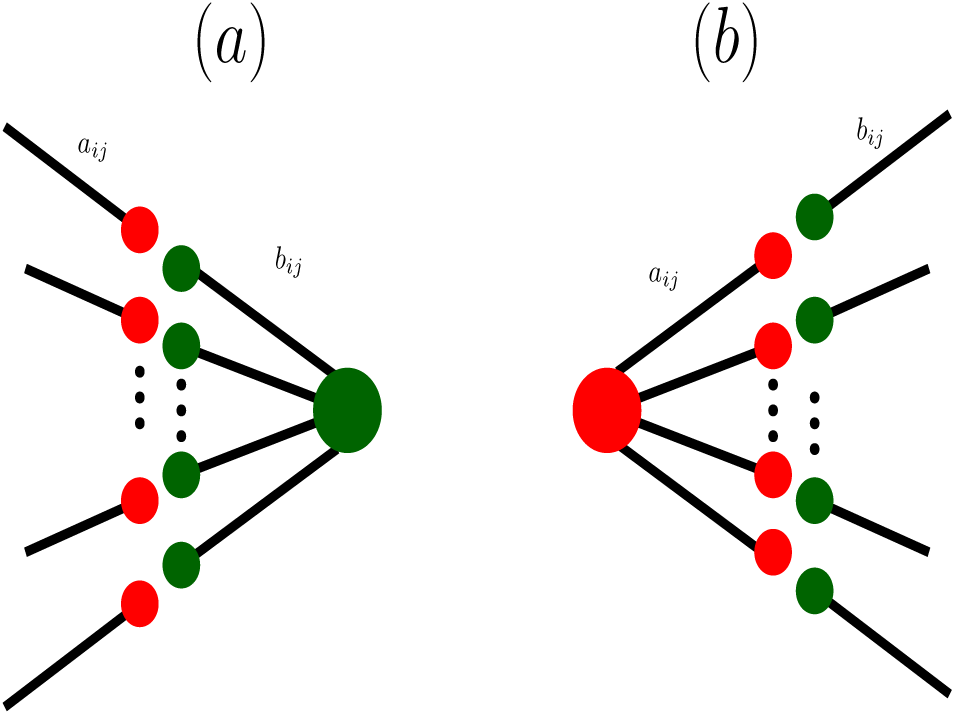
Illustration of synaptic competition induced by hetero-synaptic plasticity. a: Competition induced by post-synaptic hetero-synaptic plasticity. Excitatory pre-synaptic neurons i target a post-synaptic neuron j. We assume that the resources required for increasing the post-synaptic components *b*_*i j*_ of the weights *w*_*ji*_ = *a*_*i j*_*b*_*i j*_ are limited and therefore distributed in a competitive manner. That is, we assume hetero-synaptic plasticity where afferent synapses that receive large eligibility signals *ε*_*i j*_ increase their efficacy while synapses that receive weaker (but for excitatory synapses always positive) signals will instead weaken. b: Pre-synaptically induced competition is assumed to follow the same principle. The signals for the changes of the pre-synaptic components *a*_*i j*_, however, are assumed to depend on the realized amount of potentiation at the post-synaptic side, i.e., to take the post-synaptic hetero-synaptic competition into account. We found that this choice is more parameter tolerant and yields more robust memory than a symmetric version where pre- and post-synaptic hetero-synaptic plasticity are based on the same eligibility signal which, however, can also realize self-organized spike pattern detection (not shown).

Here, Dale’s law is imposed: when *δa* and *δb* would change the sign of *a* and *b*, respectively, we set them to zero. We assume that correlations between pre-synaptic input and post-synaptic membrane potential drive weight changes. By considering only the spike at time 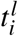 the correlation between *i*′ *th* afferent and the *j*′*th* post-synaptic neuron’s potential *V*_*j*_(*t*) is:

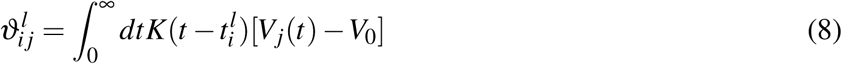

where *V*_0_ is the modification threshold^33^ that is set to zero in all simulations of this study. In every learning epoch, all *n*_*i*_ spikes in the *i*′*th* afferent are used for updating the weights. As a result, we compute correlation for all spikes *l* and add them together to determine the eligibility

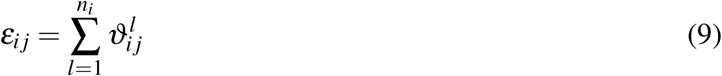

which is the basic signal for weight changes at synapse (*i, j*).

For inhibitory synapses weight changes are set to be simply proportional to *ε*. For excitatory synapses changes are assumed to depend only on the positive part [*ε*_*i j*_]_+_ which mimics the characteristics of the NMDA receptor ^16–18^. For excitatory input we also take synaptic scaling into account. That is, neurons tend to fire and to implement a specific firing rate *r*_0_ that is genetically determined ^19^. Therefore, if a post-synaptic neuron’s long-term firing rate *r* _*j*_ is less than the desired *r*_0_, it scales its afferent excitatory weights up. When *r* _*j*_ is larger than *r*_0_, it does not provide resources for further growth. We implement this with a quantity *q* _*j*_:

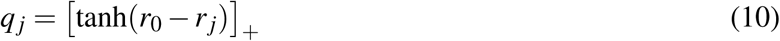

Changes of synapses are subject to limitations of the material provided by the respective pre and post-synaptic neurons (e.g., release sites, vesicles, receptor densities). Plausibly, this leads to competition for changes of different synapses, which affects the respective pre- and post-synaptic components a and b differently. In the current approach we include pre- and post-synaptic competition only for the changes of excitatory weights thereby modeling pre- and post-synaptic versions of hetero-synaptic plasticity ^20, 25, 34, 35^.

#### Single post-synaptic neuron

To compute weight change (Eq. 7) when there is only a single post-synaptic neuron (*j* = 1), we have *b*_*i*1_ =: *b*_*i*_, *ε*_*i*1_ =: *ε*_*i*_, *q*_1_ =: *q*, and *a*_*i*1_ =: *a*_*i*_. For simplicity we here set *a*_*i*_ = 1 and *δa*_*i j*_ = 0 (for the full version see next section).

Here inhibitory synapses are changed by

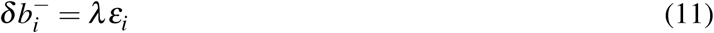

where *λ* is the learning rate for inhibitory synapses.

For excitatory synapses we subtract the mean of the plasticity signals [*ε*_*i*_]_+_, to mimick post-synaptic hetero-synaptic plasticity:

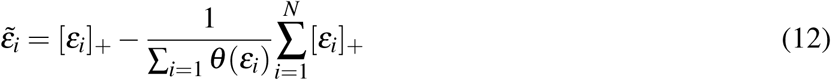

where *θ* is the Heaviside step function. Note, this does not imply strict normalization since Dale’s law prevents negative changes that otherwise would turn excitatory synapses to inhibitory synapses.

The excitatory afferents’ weights then change according to the following equation:

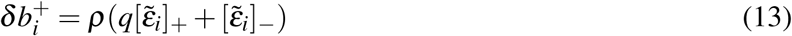

where *ρ* is the learning rate.

#### More than one post-synaptic neuron

When there is more than one post-synaptic neuron each afferent has different eligibility for each post-synaptic neuron (Eq. 9). Therefore *δb*_*i j*_ needs to be taken into account (Eq.7).

For inhibitory synapses we use:

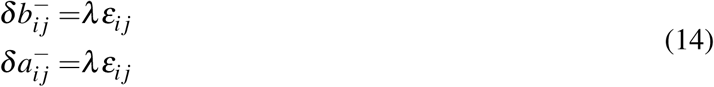

As for single target neurons *j* the changes of excitatory synapses are subject to post-synaptic hetero-synaptic plasticity that we realize by subtracting the mean of eligibilities:

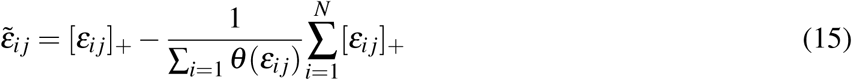

With this the post-synaptic components change by

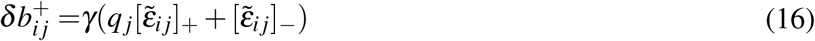

where *γ* is the learning rate. We assume that also pre-synaptic neurons have finite resources for increasing their contributions to synaptic efficacies (*a*_*i j*_), as e.g., release sites or vesicle densities. Fig 9 illustrates this pre- and post-synaptic competition. We restrict the competition to excitatory neurons that up-regulate synapses because this requires resources, whereas decreases of efficacies may even release resources. To apply the competition, we first identify the signals that would scale up excitatory synapses:

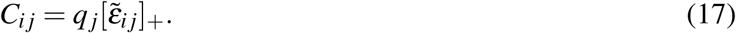

The pre-synaptic competition then is implemented by subtraction of the mean of *C*_*i j*_

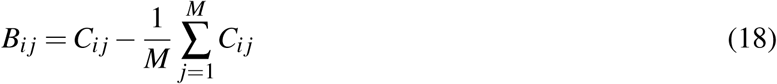

where M is the number of post-synaptic neurons. Therefore

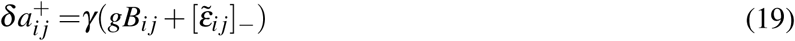

where *g* is a constant to accelerate pre-synaptic competitions.

#### Simulation details

In all simulations, there are 500 afferents (80% excitatory and 20 % inhibitory), and the learning epoch length is 2000 ms except for Fig 6 (800 ms). Afferent and embedded pattern spikes are generated randomly with Poisson point processes in which the excitatory rate (*r*_*e*_) is 5 Hz, and the inhibitory rate (*r*_*i*_) is 20 Hz. Except for Fig 8 (blue line), all embedded patterns are in the epoch in every learning cycle.

In the network model, the initial synaptic efficacies of *a*_*i j*_ and *b*_*i j*_ are chosen from a Gaussian distribution with a mean of 0.1 and standard deviations of 0.1 (negative numbers are set to zero). In the single post-synaptic neuron model, they are chosen from a Gaussian distribution with a mean of 10^−2^ (0.1 for Fig 6) and standard deviations of 10^−3^. (negative numbers are set to zero).

Synaptic scaling depends on firing rates averaged over long times. Since we perform on-line learning we use a low pass filter such that the rate estimation *r* _*j*_(*c* + 1) of post-synaptic neuron *j* used for learning in epoch *c* + 1 is a running average of the actual rates 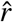 in previous epochs:

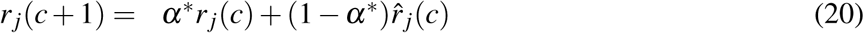

The parameter *α*\* is 0.9 for all figures. Furthermore, plasticity is not instantaneous but depends on an accumulation of signals for weight changes over some time. We implement also this by a low pass filter such that the changes taking place in each epoch 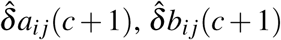 are running averages depending on the previous raw change signals *δa*_*i j*_(*c*), *δb*_*i j*_(*c*)

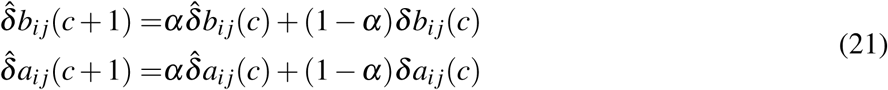

Initial conditions are 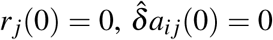, and 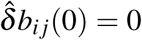. The parameter *α* is 0.99 for all figures except Fig 6 in which *α* is 0.9, *ρ* = 1.8 *×* 10^−4^, and *λ* = 2 *×* 10^−4^. Because synaptic strength cannot become arbitrarily strong due to synapses’ structure and other constraints, we cut weights changes that would carry those synapses out of the bounds which are plus and minus one for excitatory and inhibitory synapses, respectively.

Besides, Dale’s rule dictates that excitatory and inhibitory synapses cannot turn into each other; therefore, we assume synaptic weights become zero if weight changes would turn their kind during the learning. As a result, subtracting the mean value in equations 12 and 18 will not change a synapse’s type (equation 7).

## Acknowledgments

This work was supported by DFG - grant (PA 569/5-1). We would like to thank Udo Ernst and David Rotermund for fruitful discussions and comments.

